## Supplemental material for "A novel SNF2 ATPase complex in *Trypanosoma brucei* with a role in H2A.Z-mediated chromatin remodelling"

| Co-IP | gene ID | annotation | identified domains | Phyre2 modelling | NES | ident. in Co-IP No. |
|---|---|---|---|---|---|---|
| No. 1 | Tb927.11.11530 | histone acetyltransferase HAT2 | SAS2 superfamily | C-terminus: TIP60 (Cov. 67% Conf. 100%) | 5.35 | 1+2 |
| No. 2 | Tb927.11.10070 | Bromodomain, putative, Bdf3 | Bromo-domain | - | 2.3 | 1+2 |
| | Tb927.3.4140 | hypothetical protein | | btb domain (Cov. 7%; Conf. 73.3) | 1.55 | 1 |
| | Tb927.4.2340 | hypothetical protein | - | - bromo-domain from *Leishmania donovani* complexed with bromosporine (Cov. 20%; Conf. 97%); - crystal structure of tcbdf5 (Cov. 17%; Conf. 97%) | 3.25 | 1+2 |
| | Tb927.5.3210 | small ubiquitin-related modifier | UBQ/ SUMO | - | 3.05 | 1+2 |
| | Tb927.6.1070 | hypothetical protein | - | - | 4.82 | 1+2 |
| | Tb927.7.2770 | hypothetical protein | - | - | 5.81 | 1+2 |
| | Tb927.9.13320 | hypothetical protein | | SMAD/FHA domain (Cov. 26%; Conf. 93%) | 2.66 | 1+2 |
| | Tb927.11.5230 | hypothetical protein | - | ENT-domain of *T. brucei* (Cov. 35%; Conf. 100%) | 2.32 | 1+2 |
| | Tb927.11.13400 | Bromodomain, putative, Bdf5 | Bromo-domain | - | 3.18 | 1+2 |

**Table S1 Identification of a new complex associated with HAT2**

10 proteins were identified via mass spectrometry in two Co-IP experiments. The initial Co-IP was performed with Tb927.11.11530, the reciprocal Co-IP with the protein Tb927.11.10070 was performed to confirm the Tb927.4.2000 co-IP data. Only Tb927.3.4140 could be identified in the initial but not in the reciprocal Co-IP experiment. The “Annotation” column indicates the curated annotation that was found for the corresponding accession number in the TriTyp database. The” identified domains” column displays the domains that were found by BLAST search using the NCBI database. The Phyre2 modelling column indicates proteins that were identified by homology modelling. Coverage (Cov.) indicates the coverage in percent between query and template. The confidence (Conf.) represents the relative probability in percent (from 0 to 100) that the match between query and template is a true homology. The nuclear enrichment score (NES) indicates a nuclear localization if positive. The last column shows in which of the two Co-IPs the protein could be identified.

| Co-IP | gene ID | annotation | NES | ident. in co-IP No. | identified domain | Phyre2 modelling | Yeast NuA4 subunit | Domain(s) |
|---|---|---|---|---|---|---|---|---|
| No. 2 | Tb927.1.650 | conserved protein, unknown function | 4.38 | 1+2 | - | MORF4 like (Cov. 96% Conf. 95%) | Eaf3/MORF4 | Chromodomain |
| | Tb927.7.4560 | Histone acetyltransferase 1 | 2.84 | 1+2 | Tudor-knot Chromo-like domain MYST HAT | C-terminus: TIP60 (Cov. 66% Conf. 100%) N-terminus: knotted tudor domain of Esa1 (Cov. 15% Conf. 99%) | Esa1 | Tudor-knot Chromo-like domain MYST HAT |
| | Tb927.7.5310 | YEATS family, putative | 6.59 | 1+2 | YEATS-Domain | Yaf9 /GAS41 (Cov. 14% Conf. 99%) | Yaf9 | YEATS |
| No. 1 | Tb927.9.2910 | histone acetyltransferase subunit NuA4 | 4.59 | 1+2 | Eaf6 | - | Eaf6 | Eaf6 |
| | Tb927.10.14190 | hypothetical protein, conserved | 5.72 | 1+2 | - | EPL1 (Cov. 63% Conf. 98%) | Epl1 | EpcA |
| | Tb927.1.3400 | hypothetical protein, conserved | 5.40 | 1+2 | Bromodomain | Bdf5 T.c (Cov. 37% Conf. 97%) | | |
| | Tb927.6.1240 | hypothetical protein, conserved | - | 1 | | | | |
| | Tb927.8.5320 | hypothetical protein, conserved | 2.23 | 1+2 | | - | | |
| | Tb927.10.8310 | histone acetyltransferase 3 | - | 1 | MYST HAT | | | |
| | Tb927.10.9930 | PHD-zinc-finger like domain | - | 1 | | | | |
| | Tb927.11.3430 | hypothetical protein, conserved | - | 1+2 | - | - | | |

**Table S2 Identification of a new complex associated with HAT1**

11 proteins were identified via mass spectrometry in two co-IP experiments. The initial Co-IP was performed with Tb927.9.2910, the reciprocal Co-IP with the protein Tb927.1.650 was performed to confirm the Tb927.9.2910 co-IP data. Tb927.6.1240, Tb927.10.8310 and Tb927.10.9930 could only be identified in the initial but not in the reciprocal co-IP experiments. The “Annotation” column indicates the curated annotation that was found for the corresponding accession number in the TriTyp database. Proteins labelled in green exhibit a homologue in the *S. cerevisiae* NuA4 complex. The nuclear enrichment score (NES) indicates a nuclear localization if positive. The “ident. in co-IP” column shows in which of the two Co-IPs the protein could be identified. The ”identified domains” column displays the domains that were found by BLAST search using the NCBI / Interpro database. The Phyre2 modelling column indicates proteins that were identified by homology modelling. Coverage (Cov.) indicates the coverage in percent between query and template. The confidence (Conf.) represents the relative probability in percent (from 0 to 100) that the match between query and template is a true homology. The “Yeast NuA4 subunit“ column states the NuA4 complex subunit with its corresponding domain (“domain(s)” column) to which the identified trypanosome protein is homologous to.

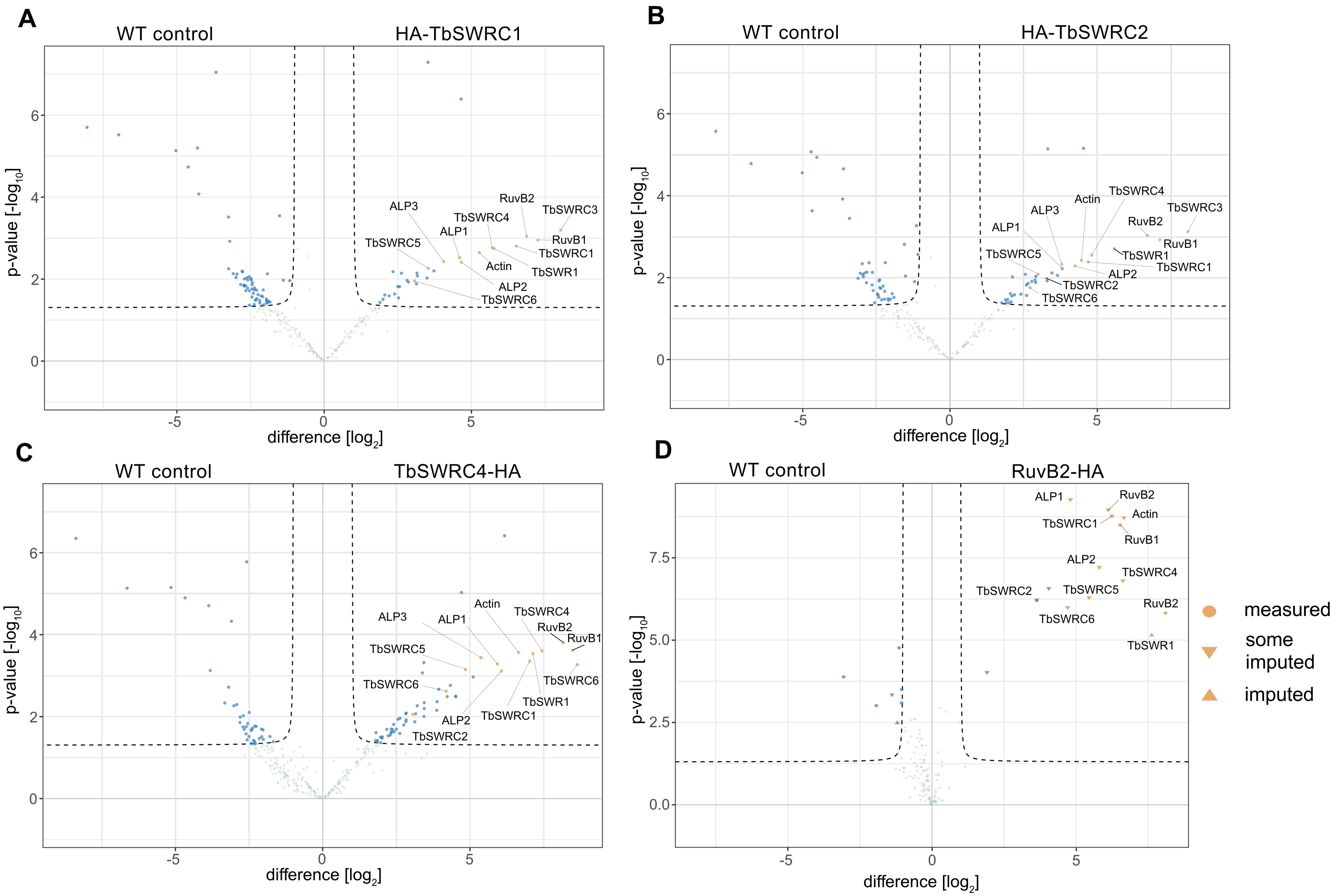
A
WT control
HA-TbSWRC1
RuvB2
TbSWRC3
ALP3
TbSWRC4
RuvB1
ALP1
TbSWRC1
TbSWRC5
Actin
TbSWR1
ALP2
TbSWRC6
B
WT control
HA-TbSWRC2
TbSWRC4
Actin
TbSWRC3
ALP3
RuvB2
ALP1
RuvB1
TbSWRC5
TbSWR1
TbSWRC1
ALP2
TbSWRC2
TbSWRC6
C
WT control
TbSWRC4-HA
Actin
TbSWRC4
ALP3
ALP1
RuvB2
RuvB1
TbSWRC5
TbSWRC6
TbSWRC6
TbSWR1
ALP2
TbSWRC1
TbSWRC2
D
WT control
RuvB2-HA
ALP1
RuvB2
Actin
TbSWRC1
RuvB1
ALP2
TbSWRC4
TbSWRC2
TbSWRC5
RuvB2
TbSWRC6
TbSWR1
p-value [-log10]
difference [log2]
measured
some imputed
imputed

**Fig. S1 Summary of volcano plots overview of 4 co-IPs**

Volcano blot of co-purified proteins after **(A)** WT control vs. HA-*Tb*SWRC1 (Tb927.10.11690), **(B)** WT control vs. HA-*Tb*SWRC2 (Tb927.11.5830), **(C)** WT control vs. *Tb*SWRC4-HA (Tb927.7.4040) and **(D)** WT control vs. HA-RuvB2 (Tb927.4.2000), co-IPs obtained by MS analysis of four biological replicates. Green dots represent purified proteins with a p-value of > 0.01 or with a fold-enrichment of = / > 1. Blue dots represent purified proteins with a p-value = / < 0.01 or with a fold-enrichment of > 1. Orange dots represent the proteins with a p-value = / < 0.01 or with a fold-enrichment of > 1 that could be identified in at least three of the four co-IPs. The annotations "measured" indicates that a sufficient number of unique peptides of the protein could be detected in the control samples to identify the corresponding protein. The annotation "some imputed" or "imputed" indicate that a theoretical value had to be imputed for some unique peptides that were used to identify the protein.

**A**

| # | Template | Alignment Coverage | 3D Model | Confidence | % i.d. | Template Information |
|---|---|---|---|---|---|---|
| 1 | c3fk3B | Alignment | 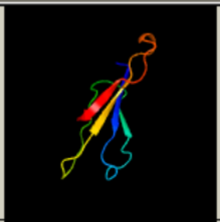 | 98.4 | 25 | **PDB header:**transcription<br>**Chain:** B: **PDB Molecule:**protein af-9 homolog;<br>**PDBTitle:** structure of the yeats domain, yaf9<br>Phyre2 Run Investigator |
| 2 | c4tmpA | Alignment | 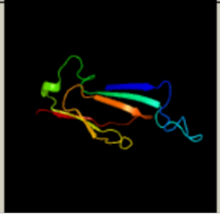 | 98.4 | 28 | **PDB header:**transcription<br>**Chain:** A: **PDB Molecule:**protein af-9;<br>**PDBTitle:** crystal structure of af9 yeats bound to h3k9ac peptide<br>Phyre2 Run Investigator |

**B**

| # | Template | Alignment Coverage | 3D Model | Confidence | % i.d. | Template Information |
|---|---|---|---|---|---|---|
| 1 | c5chlA | Alignment | 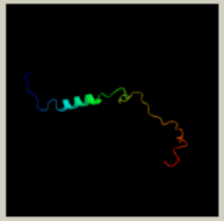 | 98.3 | 33 | **PDB header:**chaperone<br>**Chain:** A: **PDB Molecule:**vacuolar protein sorting-associated protein 72 homolog;<br>**PDBTitle:** structural basis of h2a.z recognition by yl1 histone chaperone2 component of srcap/swr1 chromatin remodeling complex<br>Phyre2 Run Investigator |

**Fig. S2 Two proteins of the new SNF2 ATPase complex exhibit features that are homologous to *Saccharomyces cerevisiae* SWR1 complex components**
**A)** Phyre2 analysis of Tb927.10.11690 (*Tb*SWRC1) exhibit an area in its N-terminus that is homologous to Yaf-9, a described component of the yeast SWR1 complex. **B)** The N-terminus of Tb927.11.5830 (*Tb*SWRC2) was identified as a YL1-domain by using the NCBI BLAST. Phyre2 analysis of the protein revealed that a part of this domain is homologous to SWC2/Vps72 which is also part of the Yeast SWR1 complex. The figure listed in the "Template" column is giving the RCSB/PDB ID of the protein that has been used for homology modelling. The "Alignment Coverage" column shows to which part of the query the template could be aligned (depicted in red). % i.d. is represents the sequence identity between query and template.

| # | Template | Alignment Coverage | 3D Model | Confidence | % i.d. | Template Information |
|---|---|---|---|---|---|---|
| 1 | c4fo0A_ | Alignment | 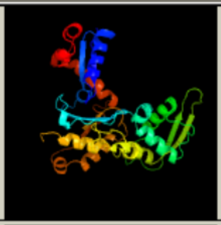 | 98.2 | 13 | **PDB header:**gene regulation<br>**Chain:** A: **PDB Molecule:**actin-related protein 8;<br>**PDBTitle:** human actin-related protein arp8 in its atp-bound state<br>Phyre² Run Investigator |
| 2 | c4i6mB_ | Alignment | 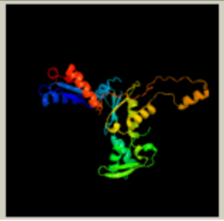 | 98.1 | 14 | **PDB header:**transcription/hydrolase<br>**Chain:** B: **PDB Molecule:**actin-like protein arp9;<br>**PDBTitle:** structure of arp7-arp9-snf2(hsa)-rtt102 subcomplex of swi/snf2 chromatin remodeler.<br>Phyre² Run Investigator |

**Fig. S3 The potential sumo interacting motif containing protein Tb927.6.2570 (*Tb*ARP2) is homologous to actin related proteins**
Homology analysis of Tb927.6.2570 revealed a homology to Arp8/9. Both Arps are essential parts of the INO80 and SWR1 complex. The figure listed in the "Template" column is giving the RCSB/PDB ID of the protein that has been used for homology modelling. The "Alignment Coverage" column shows to which part of the query the template could be aligned (depicted in red). % i.d. is represents the sequence identity between query and template.

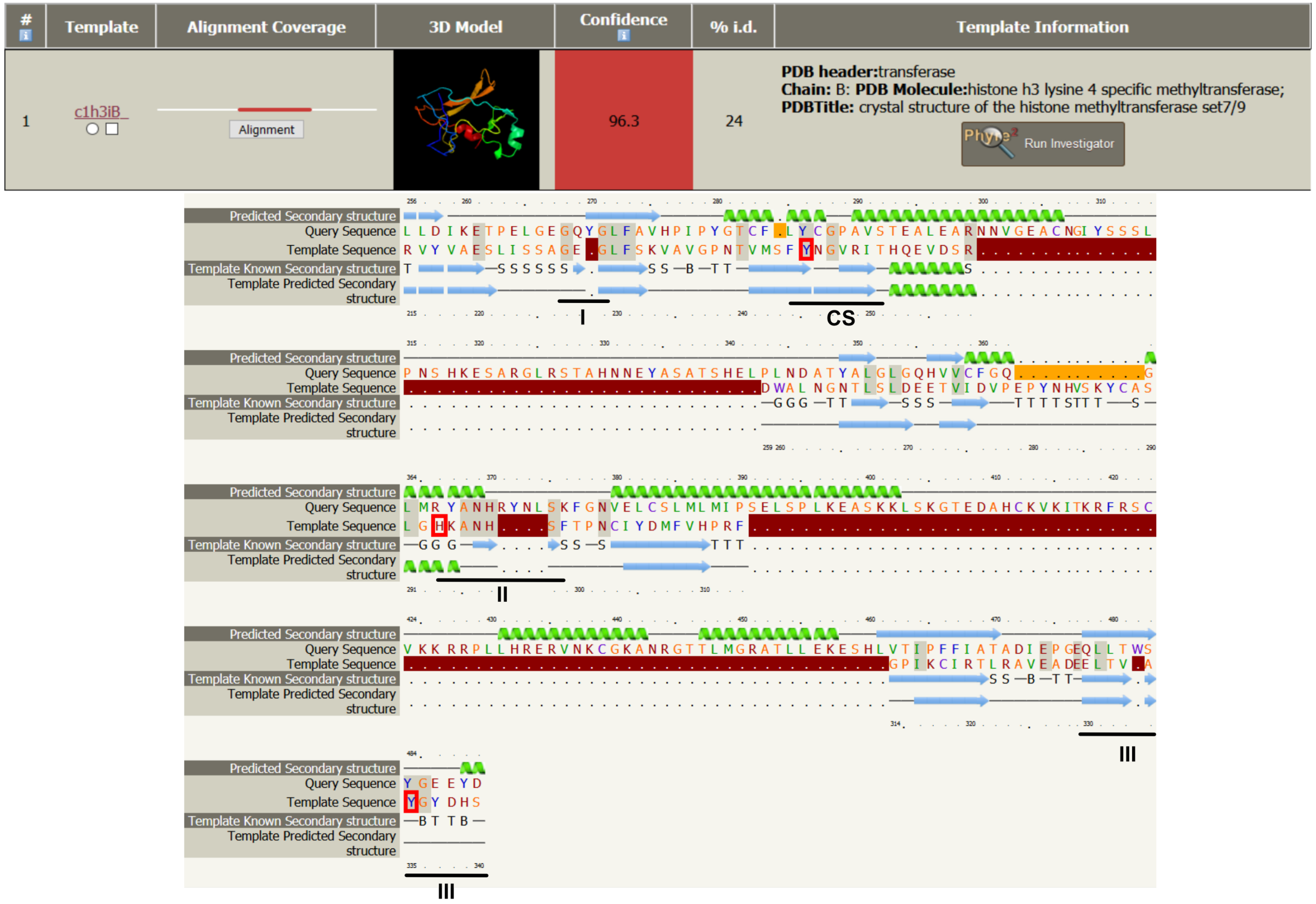


**Fig. S4 The core region of Tb927.9.8510 (*Tb*SWRC3) is homologous to SET7/9 histone methyltransferases**

Phyre2 modelling of Tb927.9.8510 could link the core of the protein to the SET7/9 methyltransferase. In the alignment of Tb927.9.8510 (Query) with SET7/9 (template) the three S-adenosylmethionin binding sites (I-III) as well as the catalytic site (CS) are highlighted. Dark-red areas represent sequence gaps in the template, yellow areas represent sequence gaps in the Query. Predicted and known β-sheets are depicted in light-blue. Predicted and known α-helices are depicted in light-green. The alignment was created by the Phyre2 online modelling platform (56). The figure listed in the "Template" column is giving the RCSB/PDB ID of the protein that has been used for homology modelling. The "Alignment Coverage" column shows to which part of the query the template could be aligned (depicted in red). % i.d. represents the sequence identity between query and template.

### *Tb*SWR1 RNAi

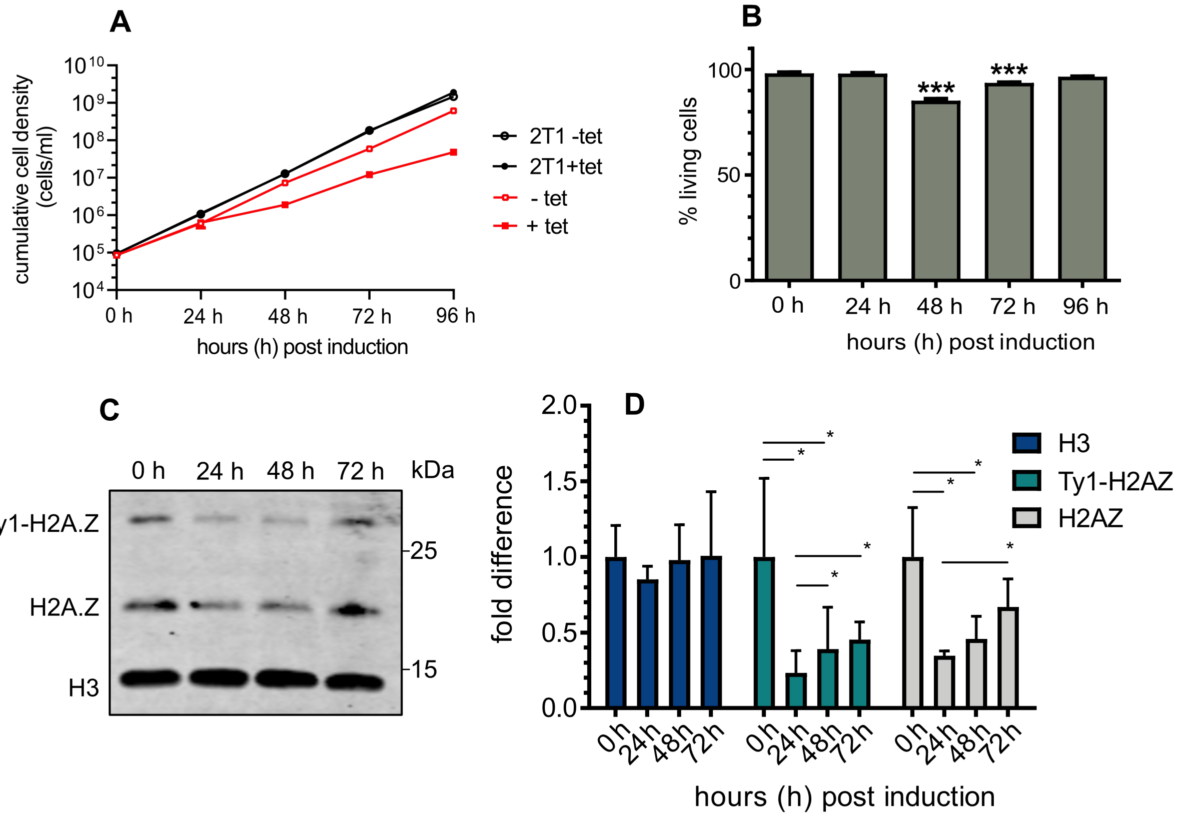


**Fig. S5 Depletion of *Tb*SWR1 reduces the amount of chromatin associated H2A.Z**
**(A)** Growth of parasites was monitored for 96 hours after RNAi-mediated depletion of *Tb*SWR1 (Tb927.11.10730) using tetraycline (tet). The parental 2T1 cell line was used as a control (n=3). **(B)** Quantification of live/dead staining with propidium iodide of *Tb*SWR1-depleted cells at the indicated timepoints post-induction. Analysis was done by flow cytometry (n=3). **(C)** Western blot analysis of the insoluble nuclear fraction with antibodies specific for histone H3 and the histone variant H2A.Z. Lysates from an equal number of cells ($2x10^6$ per lane) were analysed for each timepoint. **(D)** Quantification of chromatin-associated H3 (dark blue), Ty1-H2A.Z (turquoise) and H2AZ (grey) (N=3 for all depicted experiments; *** = p-value <0.001; ** = p-value 0.001-0.01; * = p-value 0.01-0.05).

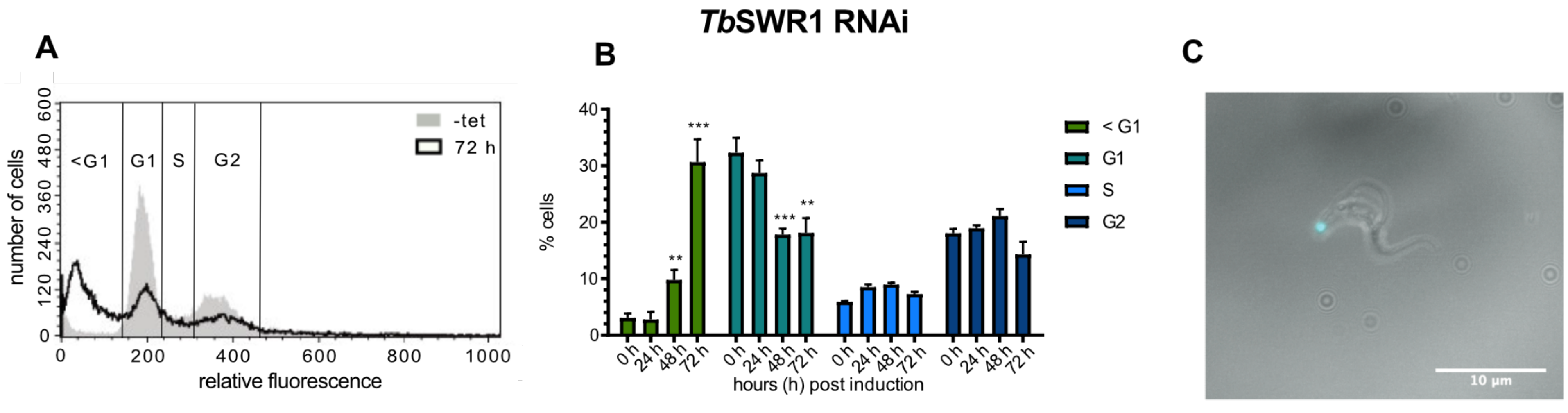


**Fig. S6 Depletion of *Tb*SWR1 leads to anucleated cell**
**(A)** Exemplary cell cycle profile of bloodstream form cells without (grey line) *Tb*SWR1 (Tb927.11.10730) depletion and after 72 h of protein depletion (black line). The Gates show the different populations of sub G1-, G1-, S- and G2-Phase cells. **(B)** Data of three triplicates, sub G1 Phase cells (green), G1-Phase cells (green-blue), S-Phase cells (light blue) and G2-Phase cells (dark blue). The data show a decrease of cells in G1 and G2 Phase in addition to the increase of sub G1-Phase cells (n=3 for all depicted experiments; *** = p-value <0.001; ** = p-value 0.001-0.01; * = p-value 0.01-0.05). **(C)** Light microscopy images (N=1) of a BSF cell after 72h of *Tb*SWR1 depletion. Scale bar 10µm.

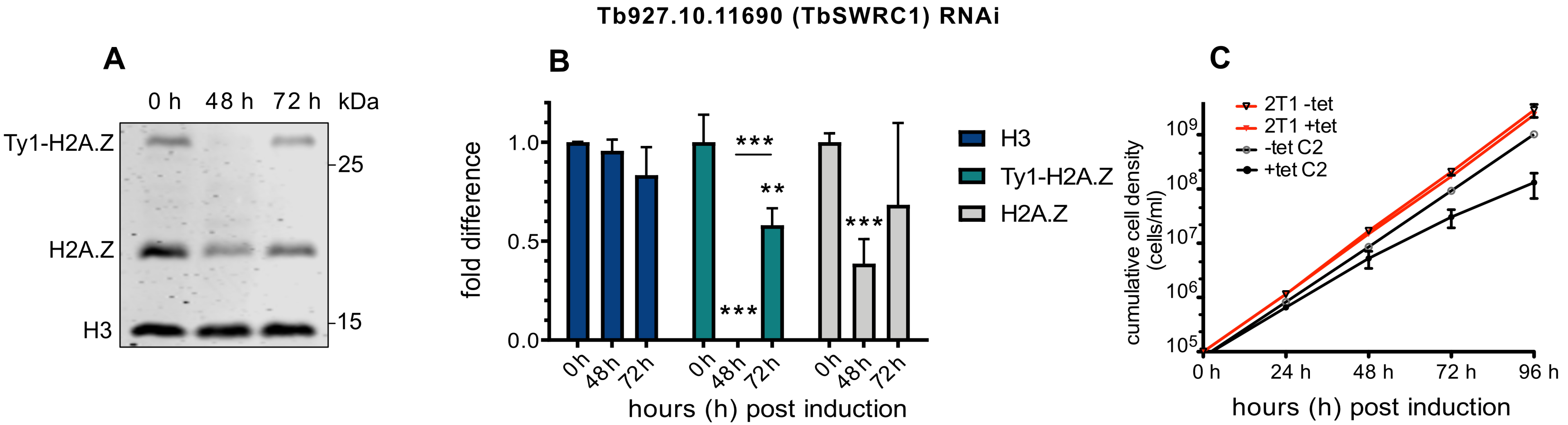


**Fig. S7 Depletion of *Tb*SWRC1 (Tb927.10.11690) reduces the amount of chromatin associated H2A.Z**
**(A)** Exemplary Western Blot analysis of the nuclear fraction with antibodies against histone H3 and the histone variant H2A.Z. An equal amount of cell equivalent was loaded for each timepoint. **(B)** The development of chromatin associated H3 (dark blue), Ty1-H2A.Z (green-blue) and H2A.Z (grey) in course of *Tb*SWRC1 depletion is plotted (N=3). **(C)** Growth of parasites was monitored for 96 hours after RNAi-mediated depletion of *Tb*SWRC1 using tetracycline (tet). Growth of tet induced and non-induced parental 2T1 cells was measured for 96h and acts as a reference (N=3 for all depicted experiments; *** = p-value <0.001; ** = p-value 0.001-0.01; * = p-value 0.01-0.05).

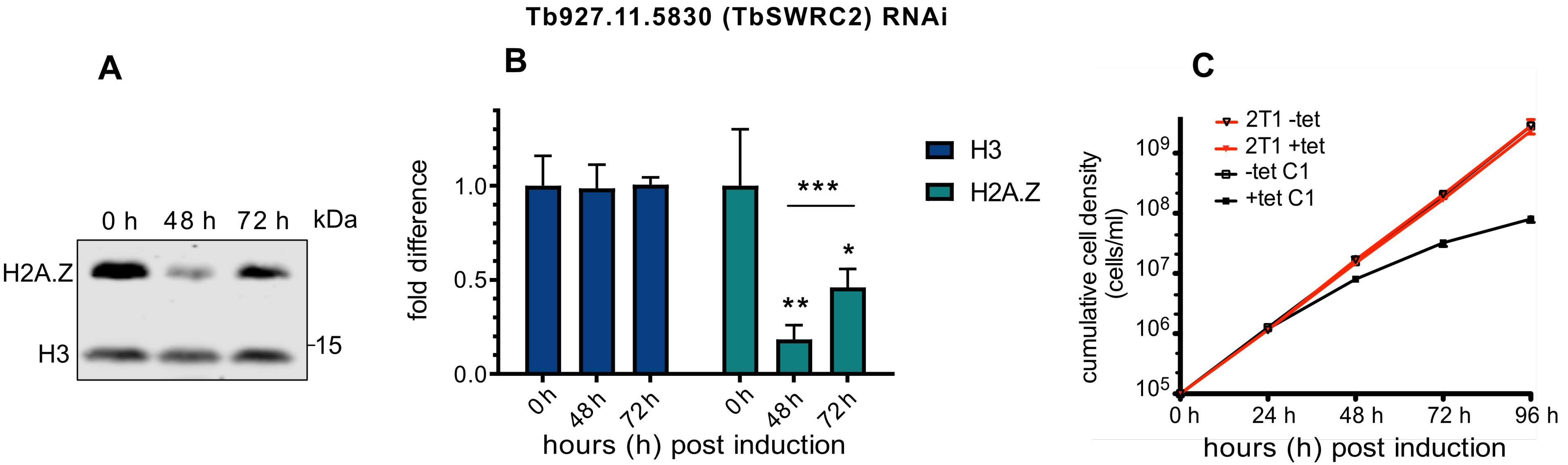


**Fig. S8 Depletion of *Tb*SWRC2 (Tb927.11.5830) reduces the amount of chromatin associated H2A.Z**
**(A)** Exemplary Western Blot analysis of the nuclear fraction with antibodies against histone H3 and the histone variant H2A.Z. An equal amount of cell equivalent was loaded for each timepoint. **(B)** The development of chromatin associated H3 (dark blue) and H2A.Z (turgoise) in course of *Tb*SWRC2 depletion is plotted (N=3). **(C)** Growth of parasites was monitored for 96 hours after RNAi-mediated depletion of *Tb*SWRC2 using tetracycline (tet). Growth of tet induced and non-induced parental 2T1 cells was measured for 96h and acts as a reference (N=3 for all depicted experiments; *** = p-value <0.001; ** = p-value 0.001-0.01; * = p-value 0.01-0.05).

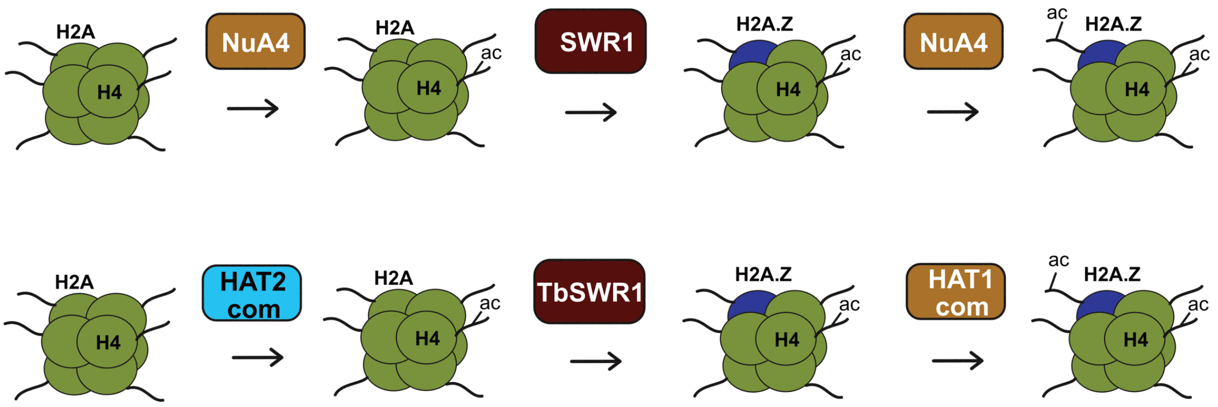


**Fig. S9 H2A.Z acetylation pathway in *S. cerevisiae* and *T. brucei***
Depiction of the H2A.Z acetylation pathway in *S. cerevisiae* and *T. brucei*. In *S. cerevisiae* the NuA4 complex acetylates histone H4 to facilitate SWR1 recruitment to the nucleosome. SWR1 exchanges H2A with H2A.Z (10, 29, 30, 58). Subsequent to the exchange the NuA4 complex acetylates H2A.Z which enhances transcription (34, 41, 42). In *T.brucei* two distinct HAT-complexes are responsible for acetylation of histone H4 and histone H2A.Z. While H4 is the substrate for the HAT2 complex, H2A.Z is acetylated by the HAT1 complex (51).

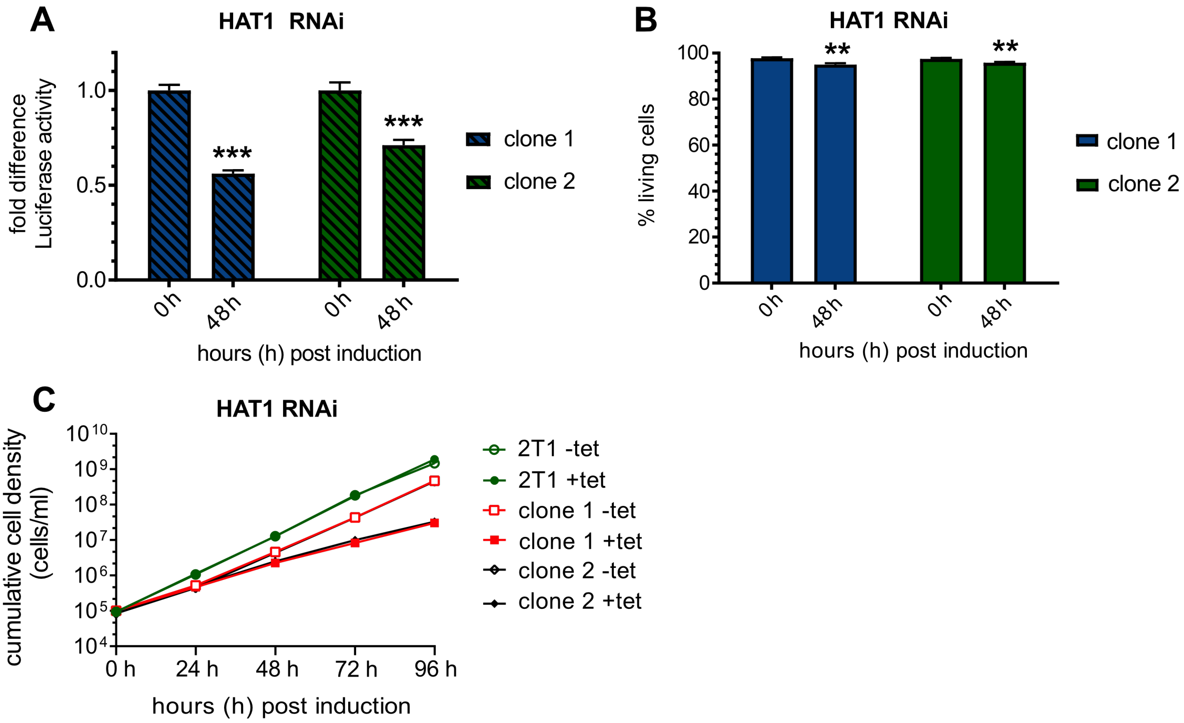


**Fig. S10 Depletion of the histone acetyltransferase HAT1 caused a decrease of reporter luciferase activity within a PTU**

A single luciferase reporter construct was integrated into the tubulin array of a HAT1 (Tb927.7.4560) RNAi cell line. Samples for the luciferase assay were normalised to cell numbers. **(A)** Luciferase activity was monitored for 48 h after induction of RNAi in two independent clones. Values of non-induced cells were set to 1. **(B)** Live/dead staining of each RNAi cell line was performed in triplicates at the same time points. **(C)** Growth of parasites was monitored for 96 hours after RNAi-mediated depletion of H2A.Z using tetracycline (tet) induction. Growth of the parental 2T1 cell line was measured for 96h as a control. (N=3 for all depicted experiments; *** = p-value <0.001; ** = p-value 0.001-0.01; * = p-value 0.01-0.05).

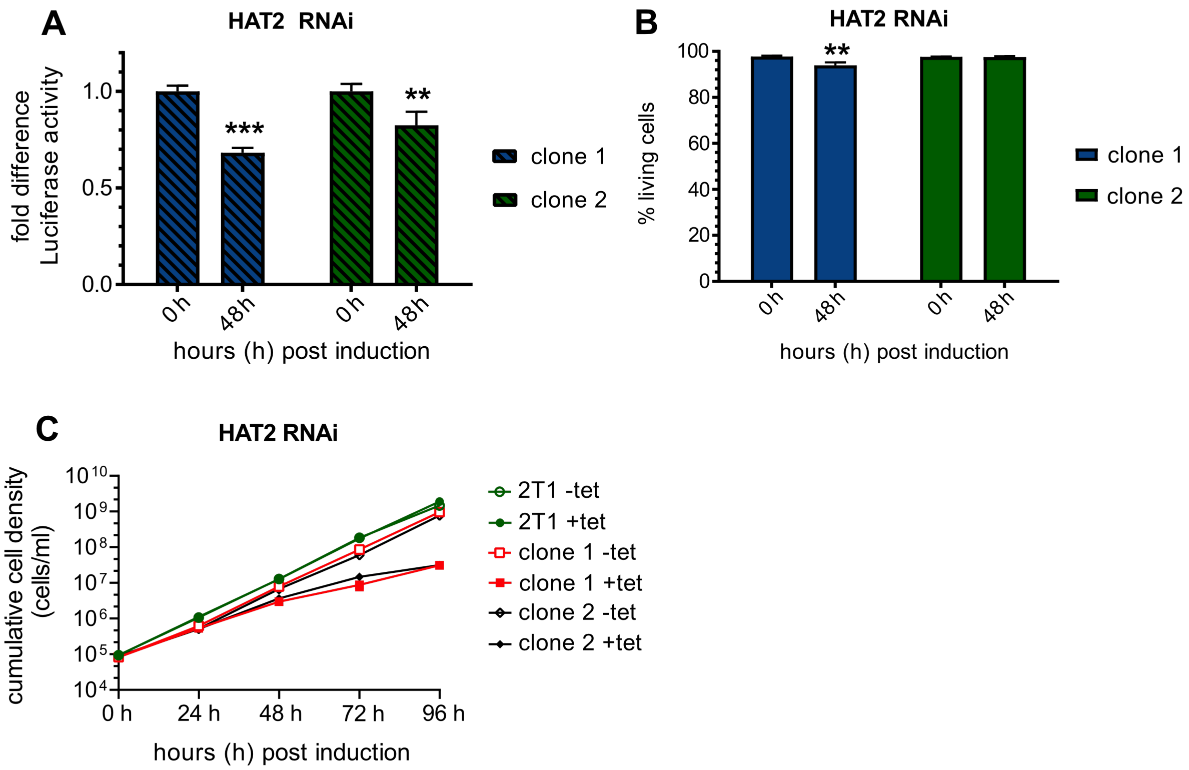


**Fig. S10 Depletion of the histone acetyltransferase HAT1 caused a decrease of reporter luciferase activity within a PTU**

A single luciferase reporter construct was integrated into the tubulin array of a HAT2 (Tb927.11.11530) RNAi cell line. Samples for the luciferase assay were normalised to cell numbers. **(A)** Luciferase activity was monitored for 48 h after induction of RNAi in two independent clones. Values of non-induced cells were set to 1. **(B)** Live/dead staining of each RNAi cell line was performed in triplicates at the same time points. **(C)** Growth of parasites was monitored for 96 hours after RNAi-mediated depletion of H2A.Z using tetracycline (tet) induction. Growth of the parental 2T1 cell line was measured for 96h as a control. (N=3 for all depicted experiments; *** = p-value <0.001; ** = p-value 0.001-0.01; * = p-value 0.01-0.05).

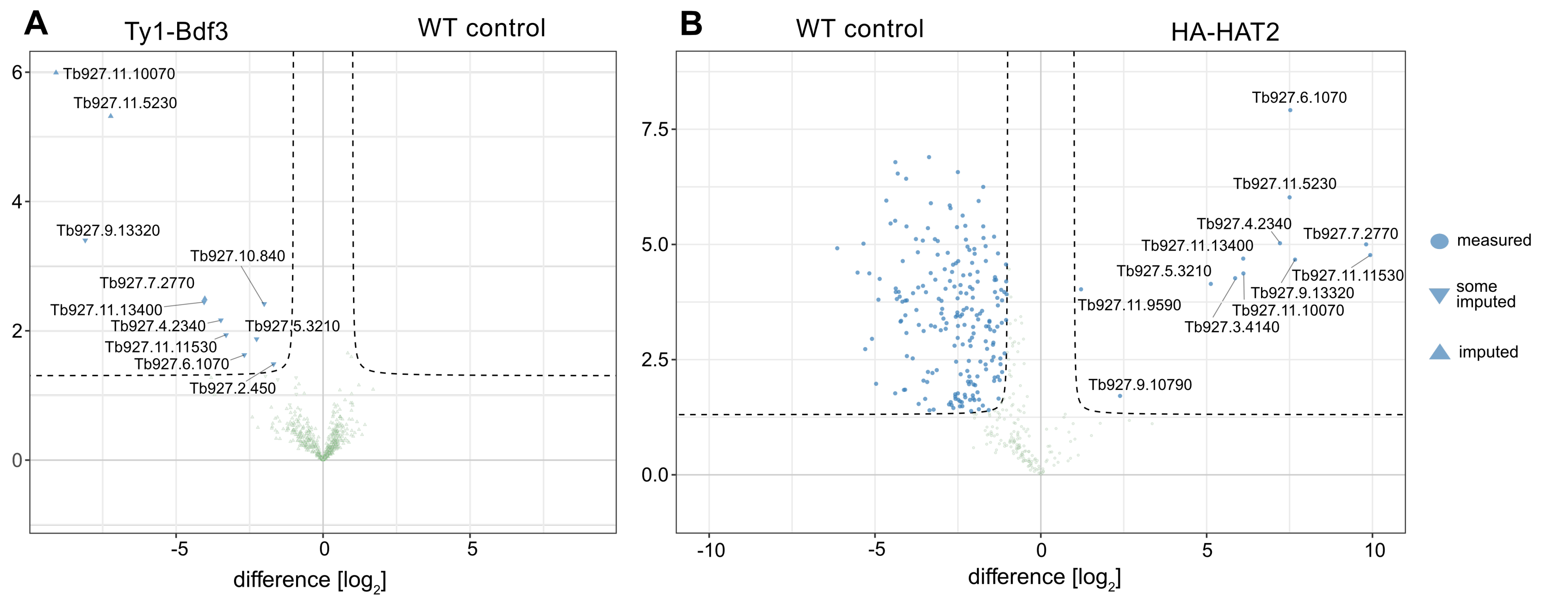


**Fig. S11 Identification of a HAT2 complex**

Volcano blot of co-purified proteins after **(A)** Ty1-Bdf3 (Tb927.11.10070) vs. WT control **(B)** WT control vs. HA-HAT2 (Tb927.11.11530), co-IPs obtained by MS analysis of four biological replicates. Green dots represent purified proteins with a p-value of > 0.01 or with a fold-enrichment of = / > 1. Blue dots represent purified proteins with a p-value = / < 0.01 or with a fold-enrichment of > 1. The annotations “measured” indicates that a sufficient number of unique peptides of the protein could be detected in the control samples to identify the corresponding protein. The annotation “some imputed” or “imputed” indicate that a theoretical value had to be imputed for some unique peptides that were used to identify the protein.

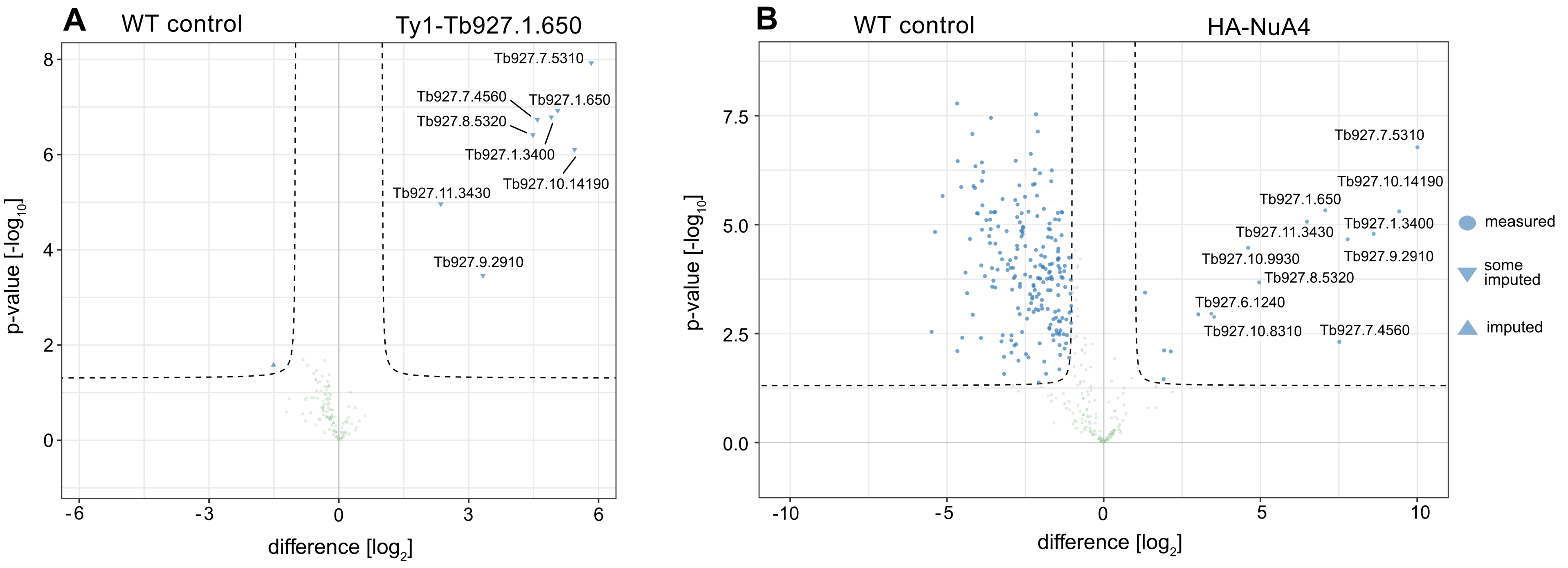


**Fig. S12 Identification of a HAT1 complex**

Volcano blot of co-purified proteins after **(A)** WT control vs. Ty1-Bdf3 (Tb927.1.650) **(B)** WT control vs. HA-HAT2 (Tb927.9.2910), co-IPs obtained by MS analysis of four biological replicates. Green dots represent purified proteins with a p-value of > 0.01 or with a fold-enrichment of = / > 1. Blue dots represent purified proteins with a p-value = / < 0.01 or with a fold-enrichment of > 1. The annotations "measured" indicates that a sufficient number of unique peptides of the protein could be detected in the control samples to identify the corresponding protein. The annotation "some imputed" or "imputed" indicate that a theoretical value had to be imputed for some unique peptides that were used to identify the protein.
